# Genome binding properties of Zic transcription factors underlie their changing functions during neuronal maturation

**DOI:** 10.1101/2024.01.04.574185

**Authors:** Melyssa Minto, Jesús Emiliano Sotelo-Fonseca, Vijyendra Ramesh, Anne E. West

## Abstract

**Background:** The Zic family of transcription factors (TFs) promote both proliferation and maturation of cerebellar granule neurons (CGNs), raising the question of how a single, constitutively expressed TF family can support distinct developmental processes. Here we use an integrative experimental and bioinformatic approach to discover the regulatory relationship between Zic TF binding and changing programs of gene transcription during CGN differentiation.

**Results:** We first established a bioinformatic pipeline to integrate Zic ChIP-seq data from the developing mouse cerebellum with other genomic datasets from the same tissue. In newborn CGNs, Zic TF binding predominates at active enhancers that are co-bound by developmentally-regulated TFs including Atoh1, whereas in mature CGNs, Zic TF binding consolidates toward promoters where it co-localizes with activity-regulated TFs. We then performed CUT&RUN-seq in differentiating CGNs to define both the time course of developmental shifts in Zic TF binding and their relationship to gene expression. Mapping Zic TF binding sites to genes using chromatin looping, we identified the set of Zic target genes that have altered expression in RNA-seq from Zic1 or Zic2 knockdown CGNs.

**Conclusion:** Our data show that Zic TFs are required for both induction and repression of distinct, developmentally regulated target genes through a mechanism that is largely independent of changes in Zic TF binding. We suggest that the differential collaboration of Zic TFs with other TF families underlies the shift in their biological functions across CGN development.

## Background

The dynamic expression and function of transcription factors (TFs) underlie the changing programs of gene expression that define stages of cellular differentiation during development (1, 2). TFs orchestrate cellular differentiation by binding in a sequence-specific manner to accessible gene regulatory elements. TFs also cooperate with co-activator and co-repressor complexes to influence the state and structure of chromatin. Thus, the regulatory function of any given TF is determined not only by when and where it is expressed, but also by a confluence of factors that determine where and how that TF is recruited to the genome (3, 4). Our understanding of the regulatory logic of TF binding has been advanced in recent years by analysis of genome-wide sequence studies that describe the chromatin and TF landscape in a wide range of different cell types and cell states (5).

Members of the zinc fingers of the cerebellum (Zic) family (Zic1-Zic5) of C2H2 zinc finger TFs are broadly expressed in dorsal neuronal progenitors during vertebrate embryogenesis (6). The Zics function to delay the exit of neural progenitors from the cell cycle, which ultimately results in the production of more neurons and larger brains (7, 8). The Zics also function in neuroblasts to promote migration, both in the embryonic brain and in the subventricular zone and rostral migratory stream of the adult rodent brain (9, 10). Knockout of the *Zic* genes in mice result in significant brain developmental defects including microcephaly, abnormal cerebellar patterning, and dysgenesis of medial structures (11). These neural progenitor phenotypes are similar to the effects of Zic loss-of-function mutations in human disorders, including cerebellar hypoplasia associated with *ZIC1* and *ZIC4* mutations in Dandy-Walker Syndrome (12), and *ZIC2* mutations in holoprosencephaly (13).

Despite their established functions as drivers of neuronal progenitor proliferation, the Zic TFs remain expressed into adulthood in select populations of differentiated neurons, including GABAergic interneurons of the olfactory bulb (10) and striatum (14), thalamic neurons (15), and most notably granule neurons of the cerebellum (16). Because *Zic* knockout mice have early developmental phenotypes, little is known about the specific functions of the Zic TFs in differentiated neurons or how they stop promoting cellular proliferation as neurons mature.

The development of CGNs in the postnatal mouse cerebellum is a useful model system to discover the mechanisms of chromatin regulation that orchestrate postmitotic stages of neuronal differentiation and maturation (17). There are temporally coordinated changes in chromatin accessibility and gene transcription that correlate with these developmental stages (18). Germline knockouts of *Zic1, Zic2, Zic3*, and *Zic4* in mice are all associated with hypoplastic cerebellum due to reduced numbers of CGNs demonstrating their requirement in CGN progenitors (6–8). In addition, Zic binding is found in the gene regulatory elements that become more accessible as CGNs mature, indicating that this TF family has functions in differentiated CGNs beyond their roles in progenitors (18). By ChIP-seq, we observed that Zic distribution across the genome changes as CGNs mature and we speculated that the shift in Zic binding could underlie a biological change in Zic function. However the functional consequences of changes in Zic TF binding for the regulation of developmental gene expression was unknown.

Here we first establish a bioinformatic pipeline to integrate Zic ChIP-seq data from the developing mouse cerebellum with other genomic datasets from the same tissue, and show how genomic location, DNA sequence, and chromatin features of Zic TF binding sites correlate with changes in gene expression over development. We then perform CUT&RUN-seq in differentiating CGNs, map Zic TF binding sites to genes using chromatin looping data and identify Zic target genes that have altered expression in RNA-seq from *Zic1* or *Zic2* knockdown CGNs. These data establish an experimentally validated set of developmentally-regulated Zic TF target genes and suggest that the collaboration of Zic TFs with other TF families defines the changing biological function of Zic TFs over the course of CGN differentiation.

## Results

### Zic TF binding consolidates from distal enhancers to promoters over CGN maturation

To characterize the genomic features of Zic binding over the course of CGN maturation we aligned Zic 1/2 ChIP-seq data (18) to the GRCm38 Gencode vM21 genome. This allowed us to compare Zic TF binding sites (peaks) to genome features and chromatin state data from other genomic datasets available from this same tissue. Of 56,941 Zic peaks, approximately 39% were significantly different between time-points (“dynamic”). 10,468 peaks were enriched at P60 (“late” peaks), and 11,721 peaks were enriched at P7 (“early” peaks). 34,752 Zic ChIP peaks were not significantly different between P7 and P60 (“static” peaks) (**Figure 1A; Additional file 2)**.

**Figure 1:**
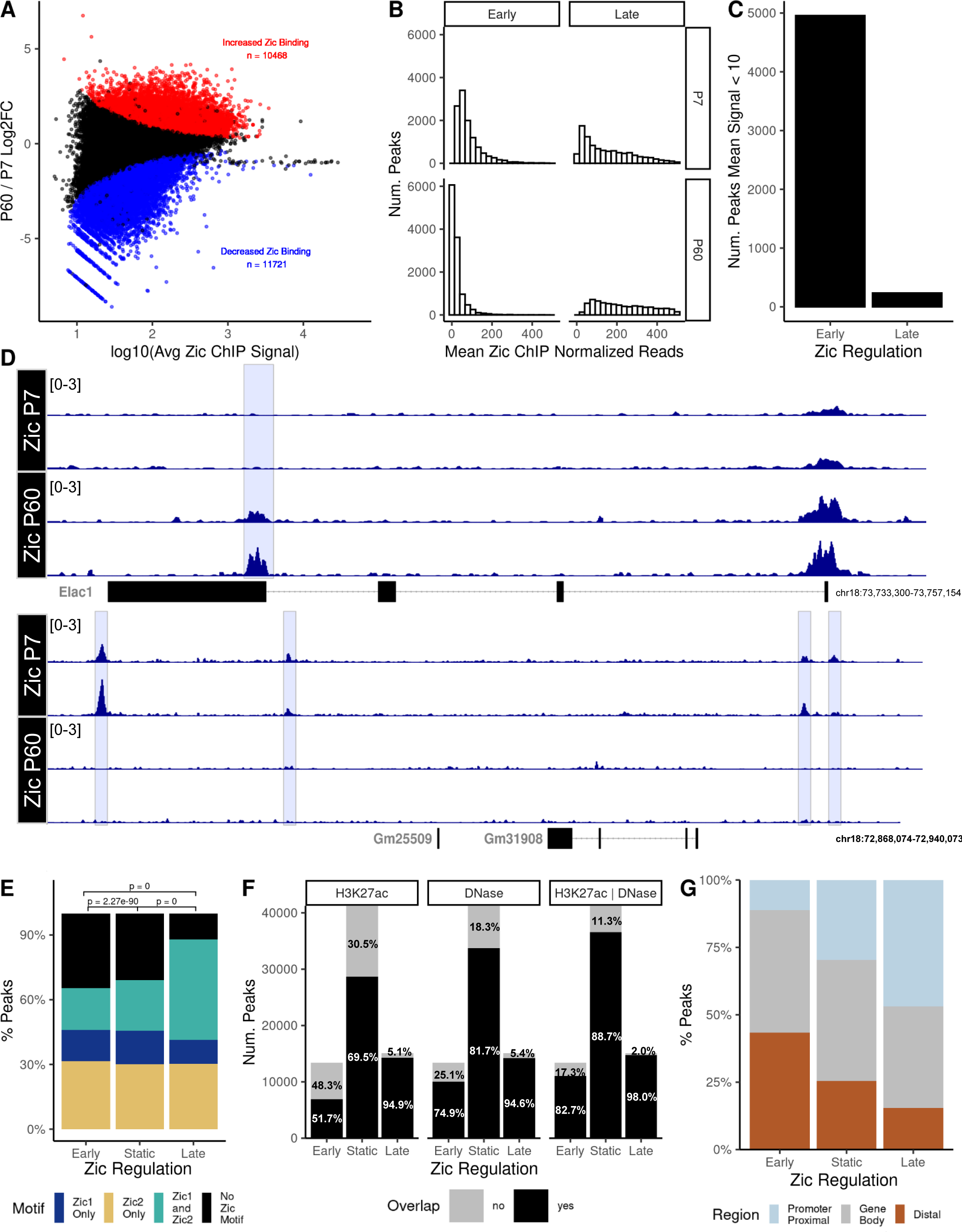
Zic1/2 binding is dynamic across mouse cerebellar development. A) MA plot comparing Zic ChIP-seq peaks at P7 and P60. Red, significantly increased, blue significantly decreased (FDR < 0.05). B) Distribution of the mean normalized reads in early and late Zic ChIP peaks at P7 and P60. C) Total number of dynamic early and late Zic ChIP-seq peaks that were either completely lost as CGNs mature (Early) or newly gained between P7 and P60 (Late) as defined in the results text. D) Example tracks of peaks that were lost as CGNs mature or gained between P7 and P60. E) Proportion of Zic1 and Zic2 motifs found in the dynamic and static Zic ChIP peaks. F) Overlap (black) or nonoverlap (gray) of Zic ChIP peaks with H3K27ac peaks, DNase hypersensitive sites (DHS), or both. G) The distribution of dynamic and static Zic ChIP-seq peaks with respect to genomic features.

Dynamic peaks could either reflect binding sites that are fully gained or lost during CGN differentiation, or they could be binding sites where the magnitude of Zic TF binding increases or decreases over time. To resolve these possibilities, we defined early and late peaks with an average read count of <10 at the other time point as those exhibiting complete loss or gain. We observed that very few (∼400) late Zic peaks had <10 normalized average reads at P7, whereas there was a much higher number of early Zic peaks (∼6000) with low average reads at P60 (**Figure 1B**, **Figure 1C**). Overall, 42.7% of the early peaks are lost as CGNs mature, whereas only 3.8% of the late peaks are newly gained (F**D**). These data show that Zic binding consolidates over time such that there is more binding at a smaller number of sites as CGNs mature.

The average width of Zic ChIP peaks is 528bp (Error! Reference source not found.**A**), which could allow for binding of multiple Zic TFs within a single peak. To assess the composition of Zic binding sites within the Zic ChIP peaks, we searched for Zic motifs in early versus late peaks. We calculated the percentage of Zic ChIP peaks that contained either Zic1 motifs (**Additional file 1: Supplemental Figure 1B**) or Zic2 motifs (**Additional file 1: Supplemental Figure 1C**) using the FIMO tool from the MEME suite (20, 34). Most peaks had only a few (0–4) Zic1 or Zic2 motifs even though the fragments were large (**Additional file 1: Supplemental Figure 1D**). Among the early and static peaks, the Zic2 motif was the most common, with a smaller proportion of peaks containing the Zic1 motif. Over 25% of peaks contained neither motif, suggesting that Zic might bind these sites in a non-canonical way either through targeting different sequences or via indirect binding. In contrast the late sites were more highly enriched for peaks with both Zic1 and Zic2 motifs (**Figure 1D-E, Additional file 1: Supplemental Figure 1D**). This supports the idea that Zic binding is consolidating at the late timepoint with the increase in both motifs and greater likelihood of direct Zic binding.

The Zic TFs are traditionally known as transcriptional activators, though in some contexts they can function in gene repression (35). Histone modifications reflect the activation state of cis-regulatory elements, with histone H3 lysine 27 acetylation (H3K27ac) serving as a marker of active enhancers and promoters. To determine whether Zic TFs are associated with active regulatory elements during early and late stages of CGN differentiation, we examined the overlap of Zic peaks with accessible and active regions of chromatin, as determined by DNase hypersensitivity (DHS) and ChIP-seq for H3K27ac at P7 and P60 (18). Early, late, and static Zic binding were all largely within regions of active chromatin indicated by overlap with DHS sites and/or H3K27ac ChIP-seq regions (**Figure 1F**). These data demonstrate that the Zic TFs are predominantly binding to open and active chromatin.

Genome-wide binding profile studies have revealed that TFs can act either by binding proximal promoters or by binding to distal enhancers, with some TFs showing a preference for one or the other location (36). Zic ChIP-seq peaks were annotated by location in the genome with respect to nearest transcription start sites, and these data showed that the distribution of Zic binding significantly shifts across CGN maturation. The early Zic peaks are nearly evenly split between gene bodies and distal enhancers, with fewer sites in proximal promoters. The late peaks are shifted in distribution toward proximal promoters, which comprise nearly 50% of all Zic peaks in the late peaks (**Figure 1G**). The static sites showed an intermediate distribution. Taken together, these data suggest that the binding of the Zic TFs consolidate from a large set of distal enhancers to a more focused set of gene promoters in maturing neurons.

### Distinct families of TFs are associated with early versus late Zic TF ChIP-seq peaks

The Zic TFs are known to cooperate with other TFs either directly through protein-protein interactions or indirectly through co-regulation of target genes (35). We reasoned that bioinformatic analysis of the Zic ChIP-seq peaks might reveal TFs that collaborate with the Zic TFs to regulate target genes. To identify these putative Zic TF co-regulators, we made the assumptions that TFs working with Zic TFs differentially over time would 1) bind close to Zic, within the regions defined as Zic ChIP-seq peaks and 2) may be differentially expressed during stages of CGN development.

We interrogated the sequence of the Zic ChIP-seq peaks to identify enriched TF binding motifs using the motif discovery program HOMER (FDR < 0.05, n = 205) (21). In parallel, we assessed the genomic locations of the early and late Zic ChIP-seq peaks for overlap with published ChIP-seq binding data for TFs using the Binding Analysis for Regulation of Transcription (BART) tool (FDR < 0.05, n = 326) (22). The combination of these methods allowed us to consider both direct and indirect genomic association of other TFs with the Zics as a possible mechanism for co-regulation of these regions (**Additional file 3**). The HOMER and BART tools each contain data on a large and overlapping set of TFs (**Additional file 1: Supplemental Figure 2A-C**). Many of the enriched TFs were shared between the early and late sites (**Additional file 1: Supplemental Figure 2A-B**). To discover TFs that may distinguish Zic function between developmental stages, we used a Rank-Rank Hypergeometric overlap (RRHO) test to find the TFs whose enrichment p-values were discordant between early and late Zic-ChIP peaks (**Additional file 1: Supplemental Figure 2D-E)**. Out of 205 enriched motifs, 35 are distinctly enriched in the early Zic peaks, and 34 and distinctly enriched in the late Zic peaks set (**Figure 2A**). Out of the 326 TFs whose ChIP binding was enriched in the early or late peak sets from BART, 53 were distinctly enriched early, and 29 were distinctly enriched late (**Figure 2B**). Distinctly enriched TFs were then filtered for concordant temporal transcriptional enrichment using the RNA-seq data resulting in 65 predicted co-regulators of Zic in early CGN maturation and 23 predicted co-regulators of Zic in late CGN maturation (**Figure 2C-D**).

**Figure 2:**
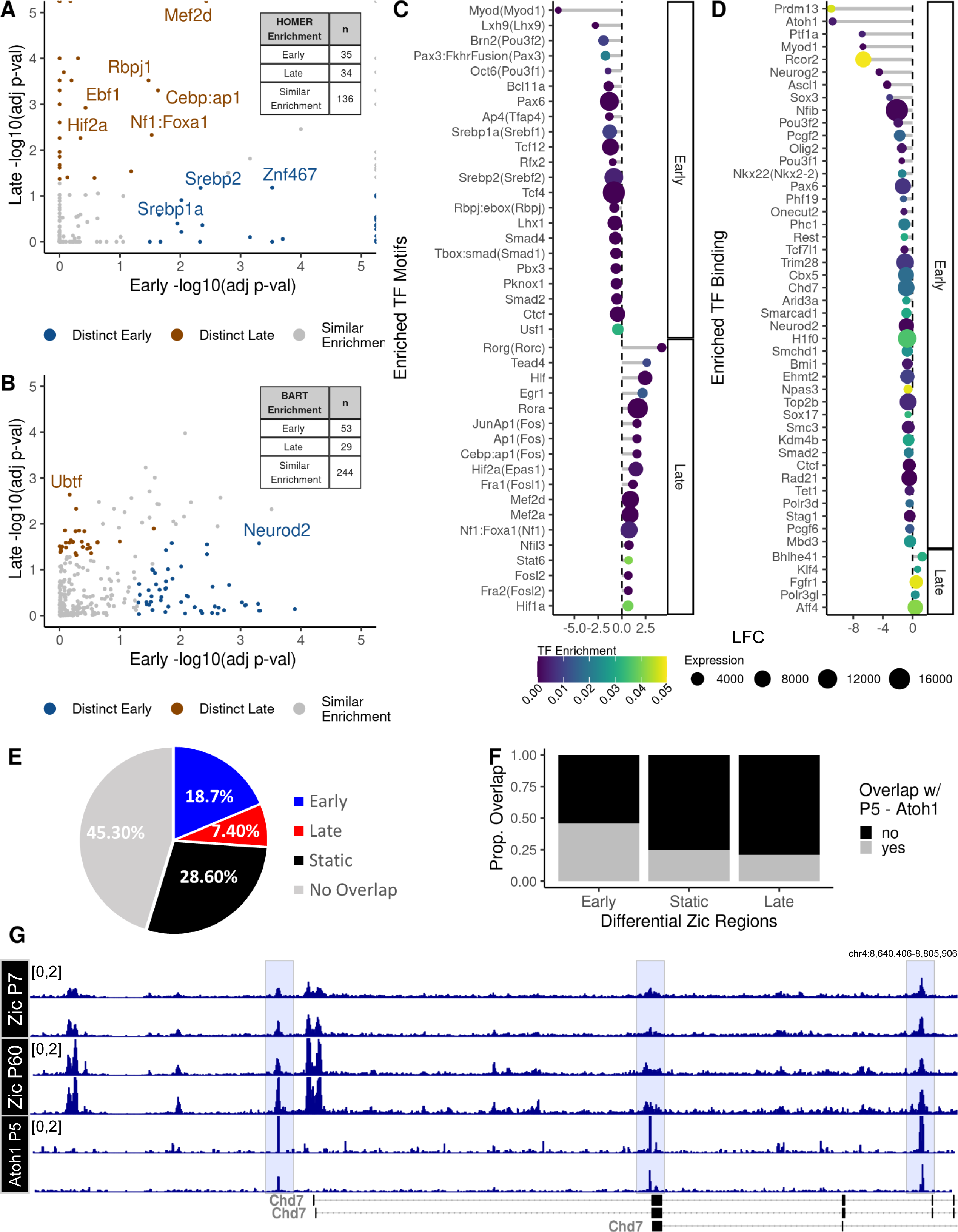
Distinct TF binding sites are enriched in early and late Zic ChIP peaks. Motif enrichment analysis using HOMER and ChIP-seq peak overlap enrichment analysis using BART was performed on the set of early and late Zic ChIP peaks to find potential collaborators of Zic TF binding. A rank-rank hyper-geometric overlap test was performed to identify the distinctly enriched A) motifs and B) TF ChIP-seq profiles between early and late Zic peaks where blue points are TF binding enriched in early Zic peaks and brown points are TF binding enriched in late Zic peaks. This set of time-point specific enriched TF C) motifs and D) TF ChIP-seq profiles within early and late Zic peaks were filtered for transcriptional enrichment at the respective time-points (P7 or P60). Each point is colored and sorted by the TF enrichment adjusted p-value, and the size of each point is the average expression of the mapped gene in RNA-seq data at the respective time point. E) The proportion of ChIP-seq peaks that are co-occupied by Zic peaks colored by the enrichment of the Zic peak (red - enriched at P60, blue - enriched at P7, black - static, and grey - no Zic peak) and F) the proportion of overlap (grey) or nonoverlap (black) of Atoh1 ChIP-seq peaks that overlap Zic peaks separated by “early” (P7 enriched), static, and “late” (P60 enriched) peaks for Atoh1 in cerebellum at P5 (25). G) Example tracks for *Chd7* at P5 overlapping with Zic binding (gray bars) in P7 or P60 cerebellum.

### Workflow captures both known and novel putative Zic co-regulatory TFs

Consistent with prior evidence that the Zic TFs collaborate with other developmental TFs in neural progenitors (37), early Zic sites were enriched for Homeobox and bHLH domain - containing TFs. Most notably, the bHLH TF Atoh1, which is a fate-determining factor for differentiation of CGN progenitors, was identified by BART as strongly enriched in the set of early Zic ChIP-seq peaks (**Figure 2D**). In the mouse cerebellum, Atoh1 is highly expressed in granule neuron progenitors from E12.5 to P14 (25, 37–40). To quantify the overlap of Atoh1 binding with the Zic TFs, we obtained a dataset of Atoh1 ChIP-seq from P5 mouse cerebellum (25) and examined the overlap of Atoh1 binding sites with our static and dynamic Zic ChIP-seq peaks (**Additional file 3**). These data revealed that 54.7% Atoh1 peaks overlap the full set of Zic ChIP-seq peaks (**Figure 2E**). Importantly, as we predicted, Atoh1 ChIP peaks overlap a greater percentage of the early Zic peaks compared with static and late Zic peak (chi-sq p-value < 0.05) (**Figure 2F**). These data showed evidence for convergent Zic/Atoh1 regulation of genes known to be important in CGN development like the chromatin regulator *Chd7* (75)(**Figure 2G**). Among the other early expressed TFs that were associated with early Zic sites were several known to be involved in cell proliferation via Wnt, FGF, Notch and SMAD signaling pathways (**Figure 2A-D**). These factors include Tfap4 (41, 42), RFX proteins (43), TCF proteins (44), which are co-effectors in Wnt/β-catenin pathways, and SMAD proteins, which are activators of TGF-beta signaling and downstream of BMP signaling (45–47). Early Zic sites are also co-localized with binding of TFs that have established functions in axon guidance (Nkx2.2) (48), and enriched for motifs of TFs that function in cellular migration (Pbx3, Pknox1, Lhx1), deepening understanding of how Zic TFs may promote CGN proliferation and migration.

Using the BART dataset in our workflow allowed us to find potential Zic co-regulatory chromatin factors. Proteins that are members of or interact with cohesin complex (CTCF, RAD21, SMCHD1, SMC3, STAG1, AND TOP2B)(49), Polycomb complexes (BMI1, PCGF2, PCGF6, PHC1, PHF19)(50), HP1 complex (CBX5, TRIM28)(51, 52), NuRD Complex (MBD3, TRIM28)(53, 54), REST complex (RCOR2, REST)(55) and BAF complex (ARID3A, BCL11A, SMARCAD1, TOP2B)(56) were all enriched in the early versus Zic binding sites (**Figure 2A,C**). We noticed that many of these are transcriptional repressor complexes or factors involved indirectly in gene regulation via chromatin architecture. These data suggest that functions of the Zic TFs extend beyond direct activation of target genes.

In contrast to our analysis of the early Zic TF peaks, we found very few hits in BART that colocalized with the late Zic TF peaks (**Figure 2D)**. This is likely a limitation of the database, which predominantly contains ChIP-seq data from dividing cells rather than postmitotic neurons. However, we did find enrichment in the late Zic TF peaks using HOMER of motifs for several TFs that show elevated expression in maturing CGNs. These include RORa and RORc, two factors involved in retinoid acid induced neuronal differentiation (57) (**Figure 2C**), as well as Hif1a which plays an important role in oxygen-dependent CGN cell-cycle exit (58) (**Figure 2D**). Most strikingly, the HOMER results suggest a role for activity-dependent transcription TFs as potential Zic TF collaborators. In the late Zic peaks, we see enrichment for binding sites of canonical activity-regulated TFs including FOSL2, FOS, JUN, EGR1, MEF2A, AND MEF2D (**Figure 2C**). At the genomic level, AP-1 transcription factors of the FOS and JUN families have been shown to promote chromatin accessibility, which can help developmentally regulated TFs to bind and could facilitate Zic binding at the late peaks we detect in mature CGNs (59). At a functional level, activity regulated TFs, especially those of the MEF2 family, are important in regulating synaptic refinement, which is a key late developmental process in postmitotic neurons (60, 61).

### Determining Zic TF regulatory activity by chromatin looping

Up to this point we have analyzed features of Zic binding with respect to their local sequence and chromatin features, but we have not yet considered the relationship between Zic TF binding and the transcriptional regulation of genes. As we show in **Figure 1K**, at most ∼50% of Zic ChIP-seq peaks are in proximal promoters, where they can be likely to regulate the nearest gene. TFs bound far away in linear space from their target genes are thought to come into close three-dimensional proximity with their target gene promoters via structural looping (62). Thus, to identify the likely target genes of the Zic TFs, and to advance understanding of the relationship between developmentally-regulated Zic binding and differential gene expression, we integrated our Zic ChIP-seq data with two different datasets of chromatin conformation (26–28) from the developing mouse cerebellum. One study used antibodies against H3K4me3 to perform promoter-centered Proximity Ligation-Assisted ChIP-seq (PLAC-seq) from adult (P56) mouse cerebellum (26) and the other used Hi-C to identify chromatin loops in cerebellum from juvenile (P22) mice (27, 28). We filtered early, late, and static Zic peaks for those that were within anchors of the captured chromatin loops in either dataset (**Additional file 1: Supplemental Figure 3, Figure 3A**). For example, the *Nr4a3* gene, whose expression increases at P60, has promoter-enhancer loops more than 600Kb upstream containing Zic peaks (**Figure 3B**). Using this approach, the intersecting Zic peaks in enhancers can be mapped to the promoters of genes they may regulate (**Additional file 4**).

**Figure 3:**
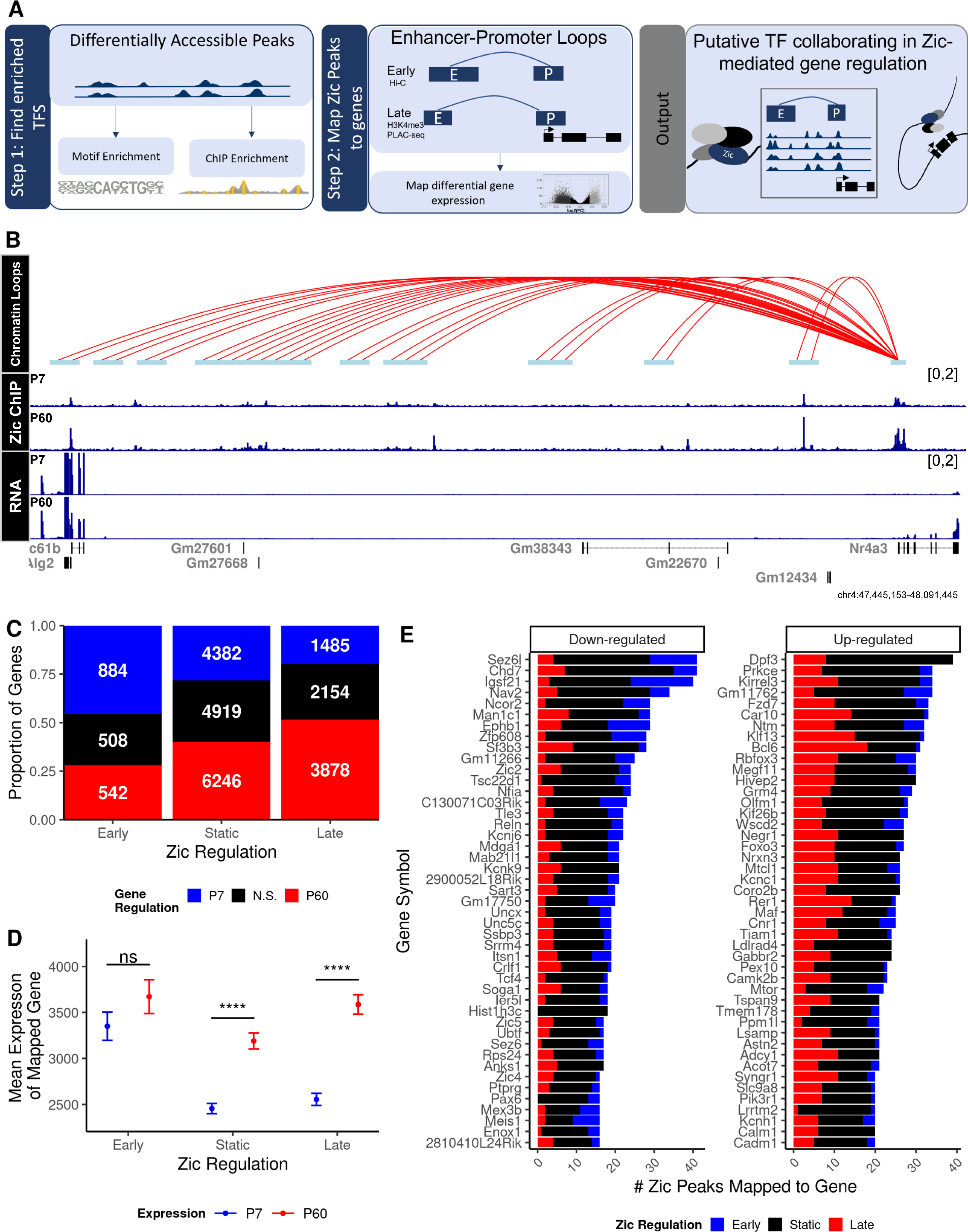
Zic binding sites can be mapped to genes through chromatin looping. Zic ChIP peaks were overlapped with anchors derived from cerebellar Hi-C (28) and H3K4me3 PLAC-seq (26) data. A) Schematic of peak mapping workflow using chromatin looping data. B) Example tracks of H3k4me3 loops interacting with the *Nr4a3* gene 100MB upstream, Zic ChIP-seq at P7 and P60, and RNA-seq at P7 and P60. C) Overall number of genes mapped to early, static, and late Zic ChIP-seq peaks. D) Expression of genes at P7 and P60 mapped to early, static, and late Zic ChIP-seq peaks. Graph shows mean and standard deviation of gene expression, *** denotes a significant difference in the mean expression between P7 and P60 with a Bonferroni adjusted p < 2.2e6 using a pairwise t-test. E) Top 50 down-regulated (FDR < 0.0f, LFC < 0) and up-regulated (FDR < 0.05, LFC > 0) genes by the number of mapped Zic ChIP-seq peaks that are dynamic between P7 and P60. Red indicates ChIP-seq peaks enriched at P60 (late), blue indicates enriched at P7 (early), and black indicates static peaks.

Though the two methods for chromatin architecture capture used at P22 and P56 differed, we expected that early Zic peaks would preferentially overlap the anchors from P22 and the late Zic peaks would preferentially overlap the anchors from P56. Indeed, a higher proportion of anchors from the early anchor dataset mapped to early Zic peaks and vice versa (**Figure 3C**). These data show that we can use chromatin conformation data to predict developmental associations of distal Zic binding sites with genes.

To determine the relationship between Zic binding and gene transcription we first assessed the average expression level at P7 and P60 of genes that map to early, static, or late Zic peaks. Overall, the expression of all the genes mapped to Zic peak-overlapping anchors rose at P60, with genes mapped to the static and late peaks showing significant increases (**Figure 3D**). Notably, the genes mapping to early Zic binding sites did not show a decrease in gene expression over time. This suggests that the loss of Zic is not a driving factor for transcriptional downregulation in maturing CGNs. However, these data do suggest that Zic has a transcriptional activating role in late stages of CGN maturation.

We next asked if the number of Zic binding events was a proxy for regulatory activity by determining if expression at any given time point or fold change in expression from P7 to P60 was a function of the number of Zic peaks that mapped to a gene. We calculated the number of early, static, and late Zic peaks that could be mapped to each gene (**Figure 3E, Additional file 1: Supplemental Figure 4**). We found a weak correlation between the number of Zic peaks and average expression (rho = 0.2, p < 2.2e16), and degree of fold change (rho = 0.11, p =1.3e14) (**Additional file 1: Supplemental Figure 4C-D**). When we looked at the top 30 genes with the most mapped Zic peaks, we saw qualitative evidence that developmentally down-regulated genes were more likely to have Zic sites that were eliminated by P60, and that developmentally induced genes were most likely to gain Zic sites (**Figure 3E**), however substantial Zic TF binding was static for both sets of genes. Taken in isolation, the number of Zic binding sites has a detectable but weak monotonic association with gene expression.

### Identification of genes that require Zic1/2 for their developmental expression

Taking chromatin looping into account helped us focus on genes that could be direct transcriptional targets of the Zic TFs. However, it is clear from our data that binding of Zic alone is not sufficient to identify genes that require Zic for their transcription; thus, incorporation of a functional molecular genomic analysis is required. In a prior study we knocked down (KD) expression of Zic1 or Zic2 in CGNs differentiating in culture and characterized changes in gene expression (18). Thus here to validate direct targets of Zic TF regulation in CGNs, we first conducted CUT&RUN-seq to map static and dynamic sites of Zic1/2 binding sites across the genome of CGNs after 1,3,5 and 7 days of differentiation *in vitro* (DIV). We then integrated these data with chromatin looping as well as RNA-seq showing dynamic changes in gene expression over this course of development in control and Zic KD neurons (**Additional file 5**).

Our CUT&RUN-seq data allowed us to refine the time course of Zic TF binding dynamics during CGN differentiation. Comparing called Zic TF peaks at DIV7 vs. DIV1 revealed 3,919 down-regulated peaks and 2,832 up-regulated peaks (FDR < 0.05), demonstrating our ability to capture dynamic Zic binding in this culture system (**Additional file 1: Supplemental Figure 5A-F**). Of the sequential comparisons, DIV3 vs. DIV1 had the greatest number of significant changes (UP = 1544, DOWN = 746) compared to DIV5 vs. DIV3 (UP = 13, DOWN = 299) and DIV5 vs. DIV7 (UP = 70, DOWN = 51). To determine how changes in Zic TF binding during differentiation in culture relate to the dynamics over the timeframe we analyzed *in vivo,* we performed a principal component analysis (PCA) of the Zic ChIP-seq and the Zic CUT&RUN peaks to cluster the samples (**Additional file 1: Supplemental Figure 5G,H**). PC1 separates the samples by developmental time, and when we considered all the samples together, the culture time points all cluster very closely together compared with the P7 to P60 separation. Along PC2, DIV1 peaks are closer to the *in vivo* P7 peaks and DIV7 peaks are closer to *in vivo* P60 peaks (**Figure 4A**). Importantly, early *in vivo* Zic ChIP-seq peaks preferentially overlapped DIV3 Zic CUT&RUN peaks and late *in vivo* Zic ChIP-seq peaks preferentially overlapped DIV7 CUT&RUN peaks (**Figure 4B**). DIV3 peaks have more overlap with early in vivo peaks (58%) than late in vivo peaks (2%) while DIV7 peaks have more overlap with late in vivo peaks (33%) than early in vivo peaks (3%) (**Figure 4B**). These data indicate that our culture results are enriched for the Zic TF binding events that happen at very early stages of CGN differentiation *in vivo* however they also support our ability to use the culture system to compare changes in Zic TF genomic binding to concordant changes in target gene transcription.

**Figure 4:**
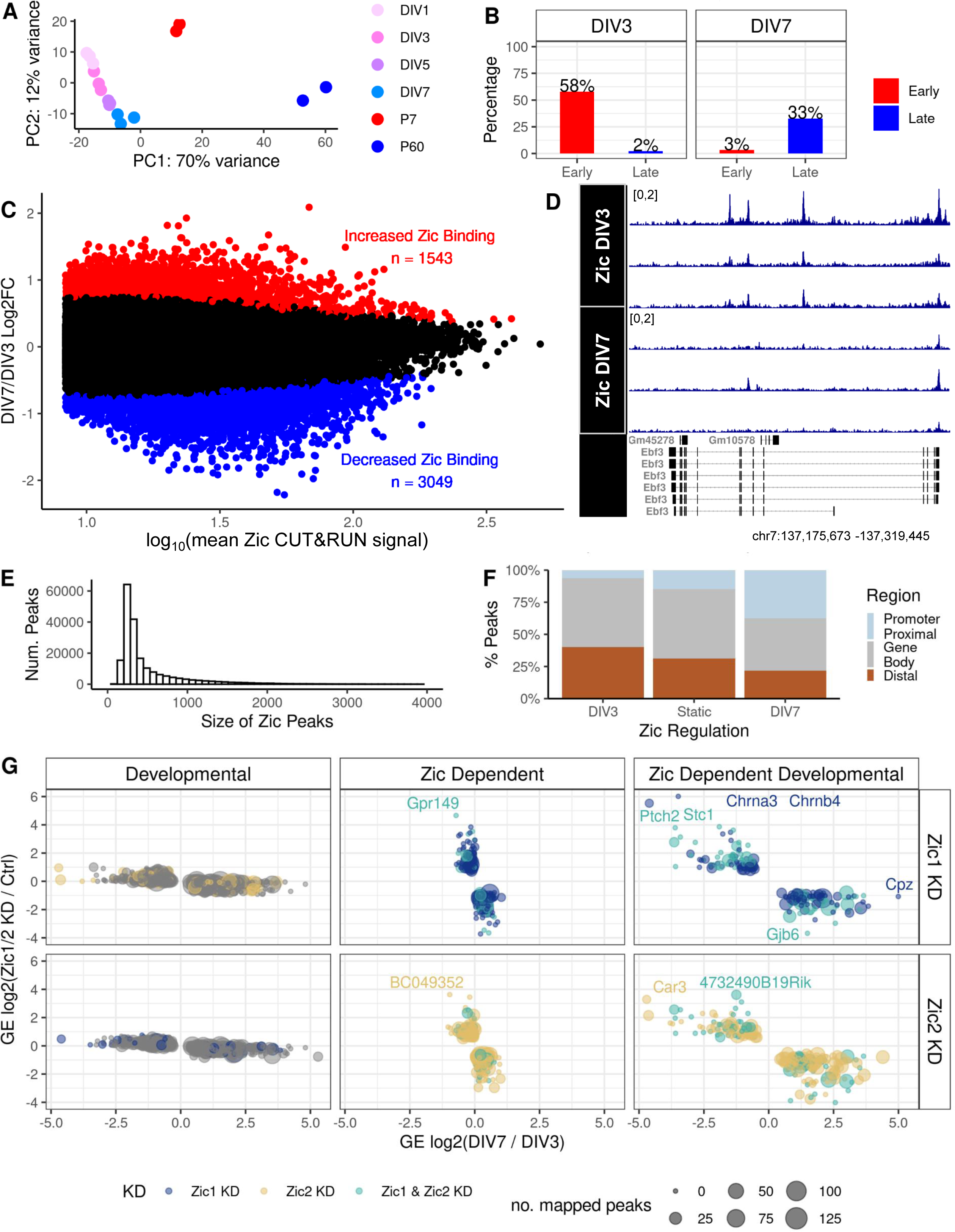
Identification of developmentally regulated and Zic-dependent genes in CGNs differentiating in culture. A) Principal component analysis of Zic binding data in culture and in vivo using the SEACR-called CUT&RUN peaks of in culture and *in vivo* samples. B) Overlap of *in vivo* Zic ChIP-seq Early and Late peaks with Zic CUT&RUN peaks enriched 3 days in vitro (DIV3) versus 7 days in vitro (DIV7). C) MA plot of Zic CUT&RUN peaks called by SEACR at DIV3 versus DIV7. D) Example of differential peak within *Ebf3* between DI3 and DIV7. E) Distribution of the size (widths) of Zic CUT&RUN peaks in a union set of the data from DIV3 and DIV7. F) The genomic distribution of Zic binding sites in DIV3-enriched, static, and DIV7-enriched Zic CUT&RUN peaks. G) Fold change of differentially regulated genes comparing DIV7/DIV3 (developmental, left) and Zic1 KD (top) or Zic 2 KD (bottom) versus shRNA control at DIV7. Genes in the left most panels are developmentally regulated genes but unaffected by Zic KD, the genes in the middle panels are significantly up- or down-regulated by Zic KD but their expression do not change from DIV3 to DIV7, and the genes in the right panels are Zic-dependent developmentally regulated genes. The colors represent whether the expression of the gene was dependent on Zic1 (dark blue), Zic2 (yellow), or both (light blue) and the size of the point represents the number of DIV3 and DIV7 union set Zic1/2 CUT&RUN peaks mapped to the gene.

We focused our analysis on the DIV3 and DIV7 Zic CUT&RUN peaks because these time points align with our previous Zic KD RNA-seq (18). Differential analysis of the 49,296 Zic CUT&RUN peaks in the merged dataset revealed 1,543 peaks enriched at DIV7, 3,049 peaks enriched at DIV3, and 44,704 static Zic peaks (**Figure 4C; Additional file 5**). One example of Zic binding loss in vitro is at the developmentally downregulated *Ebf3* gene (**Figure 4D**). Like the P7/P60 Zic ChIP-seq peaks, the DIV3/DIV7 Zic CUT&RUN peaks sizes are large enough to allow for binding of multiple TFs, with a median size of 317bp (**Figure 4E**). Additionally, the Zic CUT&RUN peaks show a similar shift from overlapping distal enhancers to consolidating at promoter proximal regions as CGNs mature (**Figure 4F**). Thus, our CGN culture system recapitulates key aspects of *in vivo* developmental Zic binding dynamics.

We used DIV3, DIV7, and Zic1/2 KD RNA-seq data (18) to identify developmentally genes that are also regulated by Zic1 and/or Zic2. Comparing DIV3 to DIV7 revealed 1388 up-regulated and 855 down-regulated developmental genes (**Additional file 1: Supplemental Figure 6A**); comparing Zic1 KD vs. control shRNA at DIV7 showed 277 up-regulated and 264 down-regulated Zic1-dependent genes (**Additional file 1: Supplemental Figure 6B**); and comparing Zic2 KD vs. control shRNA at DIV7 revealed 303 up-regulated and 435 down-regulated Zic2-dependent genes (**Additional file 1: Supplemental Figure 6C**). Finally, to identify the set of Zic-dependent developmental genes (ZDDs), we performed pairwise RRHO analyses to find the genes that showed a discordant expression upon Zic1 or Zic2 KD (**Additional file 1: Supplemental Figure 6D**). This analysis yielded genes in three categories: 1) developmentally regulated but Zic-independent (n=1582), 2) Zic1- or Zic2-dependent but not developmentally regulated (n=455), and 3) Zic1- or Zic2-dependent and developmentally regulated (ZDDs, n=329) (**Figure 4G; Additional file 5**).

We used the ZDD genes to determine how Zic binding relates to changes in gene expression. To identify direct Zic targets from the ZDD gene list, we asked which of these genes had Zic TF CUT&RUN peaks associated with their promoters via chromatin loops following the workflow described in **Figure 3A**. 37 ZDD genes had anchors that overlapped Zic CUT&RUN peaks. Notably, this analysis identified direct Zic target genes that required Zic for repression as well as genes that required Zic for induction over developmental time, which we discuss further below. If changes in Zic TF binding were driving the developmental regulation of these genes, we would predict that Zic binding events at these anchors would be dynamically regulated between DIV3 and DIV7. However, the Zic CUT&RUN peaks that mapped to the ZDD genes were mostly static between DIV3 and DIV7 (**Figure 5A**). For example, *Ets2* is a gene that fails to upregulate over time in culture when Zic1/2 are knocked down, yet most of the Zic TF binding at the *Ets2* promoter and associated enhancers is static between DIV3 and DIV7 (**Figure 5B**). Thus, similar to our analysis of differential Zic binding *in vivo,* we conclude that developmental changes in Zic binding are not required for changes in target gene expression.

**Figure 5:**
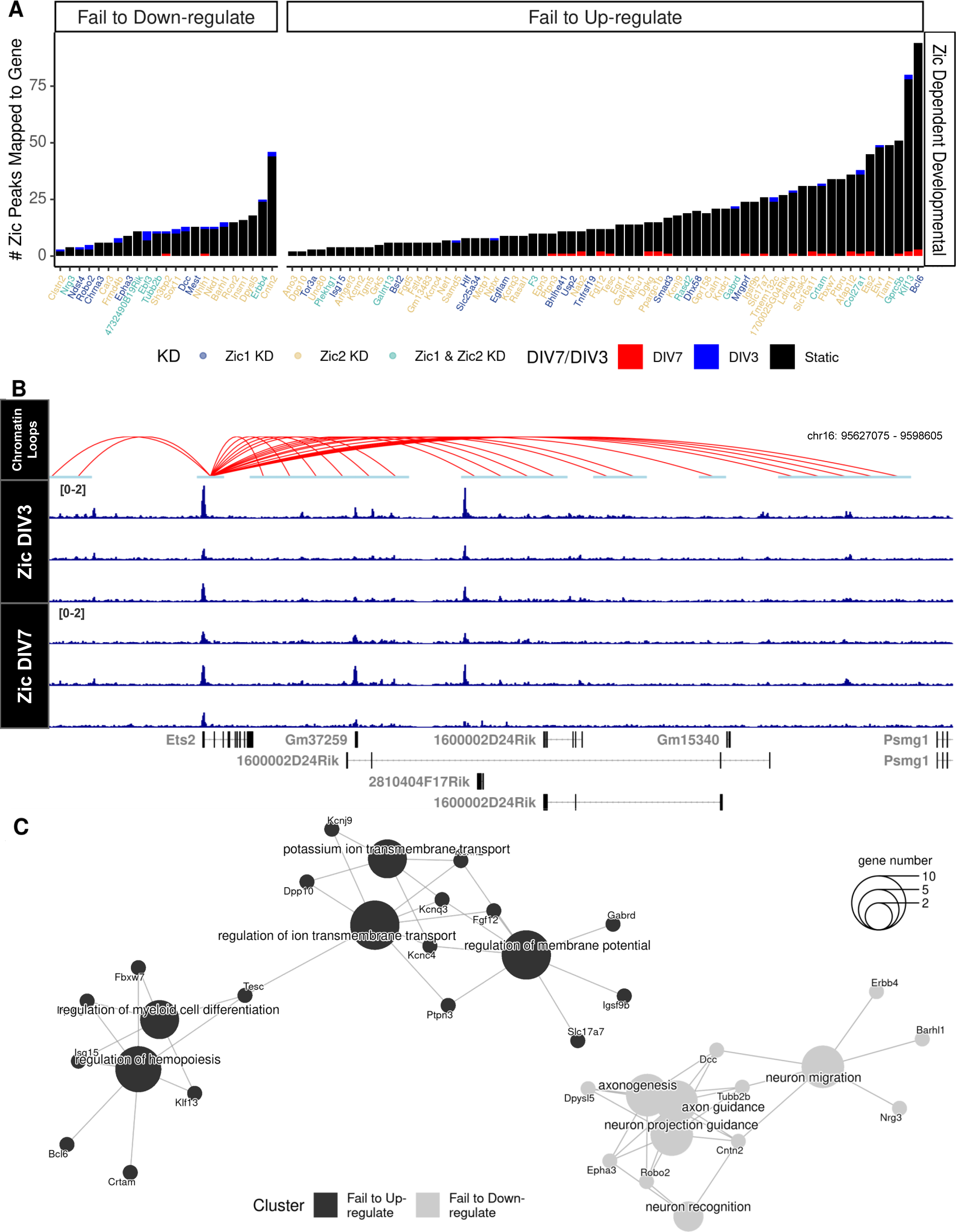
Candidate direct targets of Zic TF repression and activation converge on processes that underlie neuronal maturation. A) Zic CUT&RUN peaks were mapped to each Zic-dependent developmental gene. Colors of the bar indicate the timepoint in which peaks are enriched (Blue, DIV3 enriched, red, DIV7 enriched, black, static) and colors of the genes indicate whether the expression of the gene was dependent on Zic1 (dark blue), Zic2 (yellow), or both (light blue). Genes are separated by their developmental up- or downregulation between DIV3 to DIV7 in CGN cultures. B) Example track of static Zic TF binding with chromatin loops from cultured CGNs near a Zic-dependent gene that fails to up-regulate upon Zic KD (*Ets2*). C) Cluster diagram of biological process gene ontologies for genes that failed to be up-regulated (black) and genes that failed to be down-regulated (gray) in the Zic1 or Zic2 knockdown. The size of the center circle indicates the number of genes in each of the categories shown. The smaller circles show specific ZDD genes, and the lines connect those genes to their biological process category. Some genes are linked to more than one biological process.

Finally, because the ZDD genes are validated direct targets of transcriptional regulation by the Zic TFs and thus offer insight into the function of these TFs during CGN differentiation and maturation. To understand the functions of these Zic targets, we performed GO enrichment analysis on the Biological Processes (BP) terms for the ZDD genes (**Figure 5C**; **Additional file 5**). Zic TF target genes that were downregulated between DIV3 and DIV7 were enriched for GO BP terms including “neuron migration” and “axonogenesis” that define early stages of brain development and neuronal morphogenesis including *Dpysl5* (89), *Dcc* (63), and *Tubb2b* (64). By contrast Zic TF target genes that were upregulated between DIV3 to DIV7 were enriched for GO BP terms including “regulation of ion transport channels” and “regulation of membrane potential” that relate to aspects of neuronal function. These genes encode several synaptic receptor and ion channels including the pyruvate transporter *Slc16a11*, the GABA receptor subunit *Gabrd*, and members of the *Kcn* potassium channel and *Ptp* protein tyrosine phosphatase gene families.

## Discussion

We implemented an integrative experimental and bioinformatic approach to understand how Zic family TFs change their function over the course of CGN differentiation. By interrogating the underlying sequence and genomic context of Zic ChIP-seq peaks in early and late stages of CGN maturation, we identified developmental stage-specific features of Zic TF binding sites and determined the relationship of Zic TF binding dynamics to effects on transcription. Our results suggest that the Zic TFs both activate and repress transcription to promote maturation of postmitotic CGNs. The Zic ChIP-seq data support a model whereby Zic TFs bind widely to distal enhancers in early development to support chromatin organization and then consolidate at promotor regions in maturing CGNs to facilitate the expression of genes involved in neuronal maturation. However Zic dependent changes in developmental gene expression can occur even in the absence of changes in Zic TF binding, and we suggest that other TF families collaborate with Zic to define the regulatory logic of Zic TF function in neuronal maturation.

The Zic TFs are known to collaborate with other TFs in early development to regulate transcription in many cell types (35). For example, Zic1 has been shown to form a complex with Pax3 and Gli2 to activate the Myf5 enhancer to promote myogenesis (65). At early stages of CGN differentiation, Zic TFs co-localize at many enhancers with the basic helix-loop-helix TF Atoh1. Atoh1 is required for both CGN neurogenesis (40) and differentiation (25) and is highly expressed in CGN progenitors both in rhombic lip of the embryonic hindbrain and in the secondary proliferative zone of the postnatal external granule layer (40, 66). Unlike the constitutively expressed Zic TFs, *Atoh1* expression turns off when CGNs leave the cell cycle and migrate inward to the internal granule layer. Over the course of CGN differentiation, we observe that Zic binding is lost from about 30% of the sites where it colocalizes with Atoh1 in progenitors (**Figure 2E**). Thus, one possible explanation for the transient nature of the early Zic sites is that Atoh1 is required as a co-factor to support Zic binding at these genomic locations. A similar process has been shown to underlie maturation of motor neurons, in which persistently expressed TFs like Isl1 are handed off between a series of transient enhancers in a manner dependent on the regulated expression of fate-determining TFs like Lhx3 (67). In addition to this evidence that Atoh1 may potentially modulate Zic binding, a prior study identified Zic in a one-hybrid screen as a regulatory protein for an enhancer of *Atoh1* that is active during neural tube formation (68). We demonstrate co-localization of Atoh1 (25) and Zic binding at this *Atoh1* enhancer in early postnatal CGNs (**Supplemental Figure 7**), further supporting the co-regulatory relationship between these two TFs in early postnatal stages of CGN development.

By contrast, in P60 cerebellum, late Zic TF binding sites are enriched for sequences that can be bound by stimulus-regulated TFs of the Fos, Egr, and MEF2 families. MEF2A/D are well-described regulators of CGN synapse development and granule neuron function in motor learning (69, 70). Similar to the Zic TFs, not only do MEF2A/D bind sites that contain their canonical sequence specificity, but they also bind to regulatory elements that have AP-1 sequences, which are bound by Fos/Jun family members (70). Like the Zics TFs, MEF2A and MEF2D are constitutively expressed over the course of CGN differentiation, however these TFs are subject to stimulus-dependent regulation via post-translational modifications that can switch their functions over time (69). Phosphorylation of the Zic TFs has been shown to modifying their protein-protein interactions in other contexts (35, 53, 65, 71, 72). Whether phosphorylation of the Zic TFs changes during CGN differentiation and whether post-translational regulation of these factors contributes to differences in their function over time remain open questions.

In addition to sequence-specific DNA binding proteins, we also saw that the Zic TFs colocalize with chromatin regulators. Binding of members of the cohesin complex including CTCF, Rad21, and Smc3 were enriched at the early Zic sites, suggesting Zic TFs could contribute to the function of these complexes in establishing 3D chromatin architecture (73, 74). The CHARGE syndrome and chromodomain helicase protein Chd7 is also among the chromatin regulators co-enriched at early Zic TF binding sites. Conditional knockout of Chd7 in CGN progenitors leads to impairment of accessibility at Chd7 bound enhancers and results in an unusual pattern of cerebellar gyrification due to changes in the orientation of progenitor division (75). Zic2 is known to interact with a different chromodomain helicase, the NuRD complex factor Chd4, to maintain pluripotency in embryonic stem cells (53). Interestingly, conditional knockout of Chd4 in granule neurons leads to increased accessibility at enhancers as well as increased chromatin interactions at loop boundaries that are normally developmentally repressed (28), highlighting the potential role for chromatin conformation in Zic function (76–78).

Our culture experiments allowed us to define direct, developmentally-regulated targets of Zic TFs using a combination of CUT&RUN for Zic binding, RNA-seq in Zic1/2 knockdown CGNs, and chromatin conformation data. These data provide further support for our hypothesis that Zic TFs function both as transcriptional activators and repressors and they suggest the key biological functions that are regulated by the Zics. Although only a small set of Zic target genes require repression by Zic1 or Zic2 for their developmental downregulation, a substantial number of these genes have functions in neuronal migration and axon guidance. Migration plays an important role in between distinct stages of CGN differentiation. CGN precursors are born in the rhombic lip between E12.5-E17 in mice (40) and undergo tangential migration across the cerebellar primordium to the external granule layer where they form a secondary proliferative zone (66, 79, 80). Then upon cell-cycle exit, newborn CGNs undergo radial migration along the Bergman glia to the Internal Granule Layer (66). We found that Zic1/2 are required in cultured CGNs to turn off target genes that are critical for CGN migration (*Barhl1, Dcc, Epha3, Erbb4, Nrg3,*) (63, 81–87) and axon guidance (*Cntn2, Dpys15, Nhlh1, Tubb2b,, Robo2*) (25, 75, 80, 88–91). Interestingly, Zic2 has previously been suggested to regulate neuronal migration via its function as an activator of EphB1 and EphA4 expression in retinal ganglion cells and dorsal spinal cord neurons respectively (92–94), whereas our data show that in CGNs both Zic1 and Zic2 function as repressors of a different Ephrin ligand (*Epha3*). The consequence of deleting Zic TFs in postmitotic CGNs for cellular positioning has not studied, but our data would predict the Zic TFs might be required for the cessation of migration once newborn CGN reach the IGL, which could be studied cell-autonomously (95).

The developmentally regulated genes that are activated by Zic1/2 binding in postmitotic CGNs are overwhelmingly related to CGN maturation. The gene with the largest number of associated Zic binding sites is the transcriptional repressor Bcl6. In CGN progenitors, Bcl6 represses the expression of *Gli* genes to block sonic hedgehog-driven proliferation, which is associated with medulloblastoma (96). In cortical progenitors, Bcl6 recruits Sirt1 to the *Hes5* promoter to drive neuronal differentiation even in the presence of notch signaling, suggesting this repressor has a broad pro-neurogenic function in neural progenitors (97). A substantial group of Zic-dependent developmentally upregulated genes participate in synaptic function (*Gabrd, Slc17a7, Gprc5b*) and neuronal excitability (*Fgf12, Kcnc4, Kcnn2, Kcnq3, Kcnj9, Dpp10*), and Zic TFs appear to co-regulate groups of genes that coordinate these processes. For example, the candidate Zic TF target *Tiam1* is known to activate Rho-GTPase signal cascades to promote synaptic and dendritic plasticity (98–102). Genes that are upstream regulators of *Tiam1* (*Klf13* and Ephrins) and also those involved in Rho-GTPase signaling (*Rasal1*, *Fgd5*, *Plekhg1*, *Arhged3, Net1*) are also candidate Zic targets (103).

While this study provides substantial evidence of targets of Zic TFs during CGN development, it is important to note the limitations of these analyses. TF enrichment via BART uses published ChIP-seq data sets acquired from many tissue types and cell lines. Subsequently, binding of TFs in non-neuronal and non-CGN cell types cannot be directly inferred in this setting. To overcome this limitation, we only searched for enrichment of TFs within Zic ChIP peaks which primarily overlapped markers of open chromatin (H3K27ac peaks and DHSs) and for enriched TFs to remain in the analyses they had to be expressed at respective timepoints. Additionally, though our analyses use a combination of Zic binding and CGN gene expression from Zic1 and Zic2 KD to determine developmental targets of Zic, further studies such as CRISPR deletion of the binding sites followed RT-qPCR of candidate genes would more fully validate these targets. Finally the antibody used for ChIP recognizes both Zic1 and Zic2 but not the other Zic family members. Although Zic1 and Zic2 are the most highly expressed Zics in the cerebellum, there could be roles for Zic3-5 at some of the Zic binding sites studied here.

## Conclusions

Using a multi-omics approach we characterized the genomic features of Zic TF binding sites over stages of CGN maturation and investigated the regulatory logic of Zic TFs for gene expression during development. We show that different TF families co-bind with the Zic TFs at early versus late stages of CGN maturation and suggest that these collaborative factors shape Zic TF function. We find that Zic TFs are required for both repression and activation of gene expression as neurons mature, though these changes occur largely independent of changes in Zic TF binding. Finally we establish a validated set of direct Zic target genes in developing CGNs, which point toward functions of the Zics in migration and synaptic function.

## Methods

### ChIP-seq and DHS Data Analysis

Zic ChIP-seq, H3K27ac ChIP-seq, and DNase hypersensitivity (DHS) data from postnatal day 7 (P7) and P60 mouse cerebellum were previously generated in (18) and reanalyzed here. ChIP-seq reads were aligned to Gencode GRCm38 vM21 genome using STAR v. 2.7.2b. Duplicate ChIP reads were filtered out and peaks were called using MACS2 v. 2.1.2 with the parameters (–narrow –no-model –ext 147). bedtools2 was used to make a consensus peak set (bedtools intersect merge) and remove (bedtools subtract) the mm10 blacklisted regions (19) for differential analysis. The peak count matrix was generated by estimating the number of reads from the consensus set using RSubreads::featurecounts() v. 2.10.5. These counts were analyzed for differential enrichment between P7 and P60 using default parameters of DESeq2 v 1.36.0 (FDR adjusted p-value < 0.05).

### Zic1/2 Motif Analysis

Zic1 and Zic2 motifs were found using Find Individual Motif Occurrences (FIMO) from the MEME-Suite v. 5.3.3 (20).

### Identifying Potential TFs Co-Binding at P7 and P60 Zic ChIP peaks

A multi-pronged approach was used to predict TFs that may co-bind with Zic TFs in CGNs. First, we used a PWM-based method (HOMER v. 4.11) (21) to identify TF motifs enriched within Zic ChIP peaks, with the default random GC% matched sequence as background. Second, we used a data driven method (BART v. 2.0) (22, 23) to identify TFs that overlap with *in vivo* Zic ChIP-seq binding. We use these two methods to identify direct binding, via motif enrichment, and possible indirect binding, via enrichment of ChIP binding. In order to focus the analysis on Zic co-factors, enrichment of Zic1 and Zic2 were filtered out.

To statistically compare enriched TFs between P7 and P60 peak sets, a Rank-Rank hyper-geometric overlap test (24) was performed that compared the ranked p-value of each enriched TF at P7 to the ranked p-value of each enriched TF at P60 separately for the two methods above (BART or HOMER) as a means to calculate significantly concordant TF enrichment (hypergeometric p < 0.05). This resulted in identification of a subset of the enriched TFs in each peak set (e.g. P7) that were distinctly enriched in comparison to those from the other peak set (e.g. P60).

The gene expression for each predicted TF whose binding motif or ChIP signal was enriched within the early or late Zic ChIP peaks was calculated using previously published CGN RNA-seq data (18). To further determine TFs that play timepoint-dependent roles in CGN development, TFs were filtered for normalized mean gene expression > 100 to eliminate poorly expressed genes and for being differential expressed between P7 and P60, which was assessed by DESeq2 (FDR < 0.05, P7 vs P60).

### ChIP Overlap Analysis

The feature bedtools intersect was used to identify the early, static, and late Zic peaks that intersected with binding of the bHLH TF Atoh1 in CGNs, as determined from a previously published dataset (25). The percent of overlap was calculated by examining how many Zic ChIP peaks had at least 1bp overlap with ChIP peaks from the other dataset.

### Mapping Zic ChIP Peaks to Genes Via Chromatin Loops

Zic ChIP-seq peaks were mapped to genes using previously published predicted promoter-enhancer loops derived from adult (P56) cerebellum H3K4me3 PLAC-seq data (26) and juvenile (P22) cerebellum from Hi-C data (27, 28). ChIP peaks that overlapped the 10kb anchor bins of these loops using bedtools intersect were considered to be within the promoter-enhancer interactions. The anchors of these loops were annotated to their nearest genes using ChIPSeekR v. 1.32.1 (29). For each loop, the anchor that was nearest to a gene was deemed the promoter anchor and the other anchor was deemed the enhancer anchor. The gene mapped to the promoter anchor was assigned to the loop as the target. For cases where both anchors overlapped gene promoters, then both anchors were deemed promoter anchors and both genes were assigned to the loop.

### RNA-seq Analysis

CGN RNA-seq data were described in a previous study (18) and are reanalyzed here. Raw fastq reads were aligned to the GRCm38 Gencode vM21 genome using STAR v. 2.7.2b. Counts were extracted using HTSeq v. 0.6.1. Normalized bigwigs were made using deepTools bamcoverage v 2.0 (parameters –effectiveGenomeSize 273087177 –ignoreForNormalization chrX) and visualized using the Gviz R package v 3.15 (30). Default parameters of DESeq2 v1.36.0 was used to obtain differential expressed genes using an FDR cutoff of 0.05(31).

### CGN Cultures and Nuclear Isolation

CGNs from male and female CD1 mice at P7 were cultured following our published protocols (18). Briefly, the cerebellum was removed and dissociated with papain, granule neuron progenitors were purified by centrifugation through a Percoll gradient, and neurons were plated on poly-D-lysine coated plates in neurobasal media with B27 supplements, 1% FBS, and pen-strep. CGNs at the indicated endpoints were scraped into 1X DPBS, spun down, resuspended in Nuclei Isolation Buffer (20 mM HEPES pH 7.9, 10 mM KCl, 2 mM Spermidine, 0.1% v/v Triton X-100, 20% v/v glycerol), incubated on ice for 5 minutes, and then spun at 2,000g for 5 min at 4C. Pelleted nuclei were resuspended in Nuclei Storage Buffer (20 mM Tris-HCl pH 8.0, 75 mM NaCl, 0.5 mM EDTA, 50% v/v glycerol, 1 mM DTT, 0.1 mM PMSF) at -80C until ready to process.

### Zic CUT&RUN

CUT&RUN was performed using the CUTANA ChIC/CUT&RUN kit (EpiCypher 14-1408) as per manufacturer guidelines with the specific changes noted here. Nuclei were resuspended in Nuclei Isolation Buffer and incubated with activated ConA beads. We used an anti-Zic1/2 C-terminal antibody provided courtesy of R. Segal, Harvard Medical School (32), which is the same antibody we used in (18). CUT&RUN libraries were made using the NEB Ultra II DNA Library Prep Kit for Illumina (NEB E7645L), and NEBNext Multiplex Oligos for Illumina (96 Unique Dual Index Primer Pairs) (NEB E6440S). Library cleanup was performed prior to and after PCR amplification using 0.8X Kapa Hyperpure beads (Roche 08963851001). PCR amplification was performed with the following parameters as described in the EpiCypher CUT&RUN kit: 1) 98C, 45 sec; 2) 98C, 15 sec; 60C, 10 sec x 14 cycles; 3) 72C, 60 sec. Libraries were then pooled and 50 bp paired-end sequencing was performed at the Duke Sequencing and Analysis Core Resource on a NovaSeq 6000 S-Prime flow cell.

CUT&RUN raw fastq read files were analyzed with FastQC and processed with Trimmomatic 0.38 for quality control and adapter trimming. Trimmed reads were then aligned to the GRCm38 Gencode vM21 reference genome using STAR 2.7.2b. Duplicates were filtered from the resulting alignments with MACS2 2.1.2 filterdup keeping only one duplicate. Genome coverage was calculated using bedtools v2.25.0 genomecov. peak calling was performed with the genome coverage file using SEACR 1.3 stringent with a numeric cutoff that returned the 0.01 fraction of peaks with top signal. A union peak file was obtained with the union function from GenomicRanges 1.48.0 R package. Raw reads were counted using this union peak file as reference with the regionCounts function from the csaw 1.30.1 R package. DESeq2 1.36.0 was used to obtain differentially bound peaks between timepoints, using an adjusted p value cutoff of 0.05. Log2 fold change estimates were shrunk using the lfcShrink function from DESeq2, and the ashr method.

### Identification of Direct Gene Targets of Zic TFs in CGNs

To find genes that are both direct targets of regulation by Zic1/2 and developmentally regulated during CGN differentiation, we integrated 1) genomic Zic binding data in cultured CGNs from Cut&Run-seq with 2) chromatin conformation data to map peaks to genes, and 3) changes in the expression of those target genes over developmental time in culture in control or *Zic1/Zic2* knockdown (KD) CGNs (18). From (18) we obtained ranked lists of gene expression changes across development of CGNs (from 3 to 7 days in vitro(DIV)) in control neurons, and changes in gene expression at DIV7 comparing control with either Zic1 or Zic2 knockdown. We reanalyzed these data sets by aligning them to GRCm38 Gencode vM21 genome and performing differential expression analysis with default parameters of DESeq2 1.36.0. We first identify a set of genes that are regulated by Zic, either directly or indirectly by comparing developmentally expressed genes (DVI3 v. DIV7) to differentially expressed genes in Zic1/2 KD conditions (ZIC1/2 KD DIV& v WT DIV7) using a Rank-Rank hypergeometric overlap test (33). Here, we considered genes that were discordantly expressed between the two comparisons as Zic dependent developmental genes. We next identified genes with overlapping Zic CUT$RUN peaks in their respective promoter and enhancer anchors from the P22 chromatin looping data (28) and considered these genes to be direct targets of Zic binding. Intersecting the lists of Zic dependent developmental genes and direct Zic target genes resulted in what we called the set of direct Zic regulatory target genes. The R package clusterProfiler v 4.4.1 was used to find enriched Gene Ontology enrichments of Zic target genes, with the background set being all mouse genes.

## Abbreviations

BART: Binding Analysis for Regulation of Transcription
BP: Biological Process (reference to gene ontology enrichment)
CGN: Cerebellar Granule Neuron
CRISPR: Clustered Regularly Interspaced Short Palindromic Repeats
DIV: Days in Vitro
FIMO: Find Individual Motif Occurrences
FDR: False discovery rate
GO: Gene Ontology
HOMER: Hypergeometric Optimization of Motif EnRichment
KD: Knockdown
RRHO: Rank-Rank Hypergeometric Overlap
TF: Transcription Factor
ZDD: Zic dependent and developmental regulated

## Declarations

### Data Availability

Mouse cerebellar Zic1/2 ChIP-seq, DNase-seq, Zic1 and Zic2 knockdown RNA-seq, and RNA-seq data from CGNs at DIV3 and DIV7 or cerebellum at P7 and P60 were generated in (18) and the publicly available data can be found at GEO:GSE60731. Atoh1 ChIP-seq data are from (25) and we downloaded the data from GEO: GSE22111. Adult PLAC-seq data are from (26) and we downloaded the data from GEO:GSE127995. Juvenile Hi-C data are from (28) and we downloaded the data from GSE138822. Zic1/2 CUT&RUN data were generated in this study and can be found at GSE211309.

### Code Availability

Scripts used for this analysis can be found in this GitHub repository: https://github.com/MelyssaMinto/zic_analysis.

### Authors’ contributions

This study was conceived by M.S.M. and A.E.W. Data was gathered by V.R., analyzed by M.S.M. and J.E.S. This manuscript was written my M.S.M. and A.E.W. All authors read, contributed to editing and approved the final manuscript.

## Acknowledgements

We thank Yue Yang (Northwestern) for sharing the predicted mouse cerebellar PLAC-seq loops, Jared Goodman (Washington University, St. Louis) for sharing the predicted mouse cerebellar HI-C loops, and Irene Kaplow (Carnegie Mellon University) for critical reading of the manuscript.

