## Additional file 1 supplementary figures for "Genome binding properties of Zic transcription factors underlie their changing functions during neuronal maturation"

### Supplemental Tables

**Additional file 2**: Differential Zic ChIP-seq Peaks

**Additional file 3**: Distinctly enriched transcription factor binding from custom workflow (described in Methods)

**Additional file 4:** Chromatin Loop, anchor to gene and anchor to Zic peak mapping

**Additional file 5**: In-vitro CUT&RUN differential binding, in-vitro RNA-seq differential expression, RRHO enrichment, and GO enrichment terms for candidate Zic targets

### Supplemental Figures legends

**Supplemental Figure 1: Characteristics of Zic Peaks.** A) Distribution of the size(bp width) of all Zic ChIP peaks in the union set of all samples at P7 and P60. Consensus sequence of B) Zic1 and C) Zic2 motifs searched in the set of all Zic ChIP peaks. D) Distribution of the number of Zic1 and Zic2 motifs in early, static, and late Zic ChIP peaks.

**Supplemental Figure 2: In vivo ChIP peak overlap enrichment and motif enrichment of putative co-regulatory TFs in Zic ChIP-seq peaks**. Significantly enriched TF motifs from Homer (black) and *in vivo* ChIP peaks from BART (orange) within A) early or B) late Zic ChIP peaks are shown. TFs that are in both databases used by HOMER and BART are blue and TFs annotated with a * denotes enrichment with both HOMER and BART tools. C) Venn Diagram showing the overlap of the TFs within the BART (orange) and HOMER (black) databases. Heatmap of RRHO p-values comparing the D) Motif and E) *In vivo* ChIP enrichment of TF binding in early and late Zic peaks

**Supplemental Figure 3: Identification of Zic ChIP peaks that overlap Hi-C anchors**. Number of Zic ChIP peaks that overlap P22 (juvenile) cerebellar Hi-C anchors from [28] and P56 (adult) PLAC-seq cerebellar anchors from [26]. Data are sorted by A) the change in gene expression over time (P7 to P60) of the target gene and B) the change in Zic binding over time of the Zic peaks that overlap the anchors.

**Supplemental Figure 4: The number of Zic peaks that map to a target gene does not determine the direction of gene regulation over developmental time.** A) Count of Zic1/2 ChIP-seq peaks mapped to each gene where the color indicates whether the gene is up-regulated (red), down-regulated (blue) or constitutively expressed (black) from P7 to P60. B) Distribution densities of number of Zic peaks by expression enrichment at different developmental stages.

Scatterplots of C) mean expression and D) absolute log fold change of gene expression levels versus the number of Zic peaks.

**Supplemental Figure 5: Zic1/2 CUT&RUN time-course shows dynamic changes in binding during development of cultured CGNs.** Zic CUT&RUN was performed in cultured CGNs at 1,3, 5, and 7 days in vitro (DIV). MA plots comparing Zic peaks called by SEACR showing the differential peaks between A) DIV7 v. DIV1 B) DIV7 v. DIV3 C) DIV7 v DIV5 E) DIV3 v. DIV1 F) DIV5 v. DIV1 and G) DIV5 v. DIV3. Principal component analysis of Zic in culture and *in vivo* using the SEACR-called CUT&RUN peaks of G) All in culture samples and H) all *in vivo* samples.

**Supplemental Figure 6: Comparison of gene expression throughout a developmental time course in cultured cerebellar granule neurons**. Volcano plots show the differential genes between A) DIV7 v DIV3, B) Zic1 KD v WT at DIV7, and C) Zic2 KD v WT at DIV7. Heatmap of RRHO p-values comparing genes changing throughout WT development versus D) with Zic1 KD and E) with Zic2 KD.

**Supplemental Figure 7: Zic binding at *Atoh1* gene.** Tracks of Zic ChIP and Atoh1 ChIP binding at the *Atoh1* gene. Tracks are annotated with early, static, and late Zic ChIP peaks and show early and static Zic peaks near the *Atoh1* gene.

Supplementary Figure 1

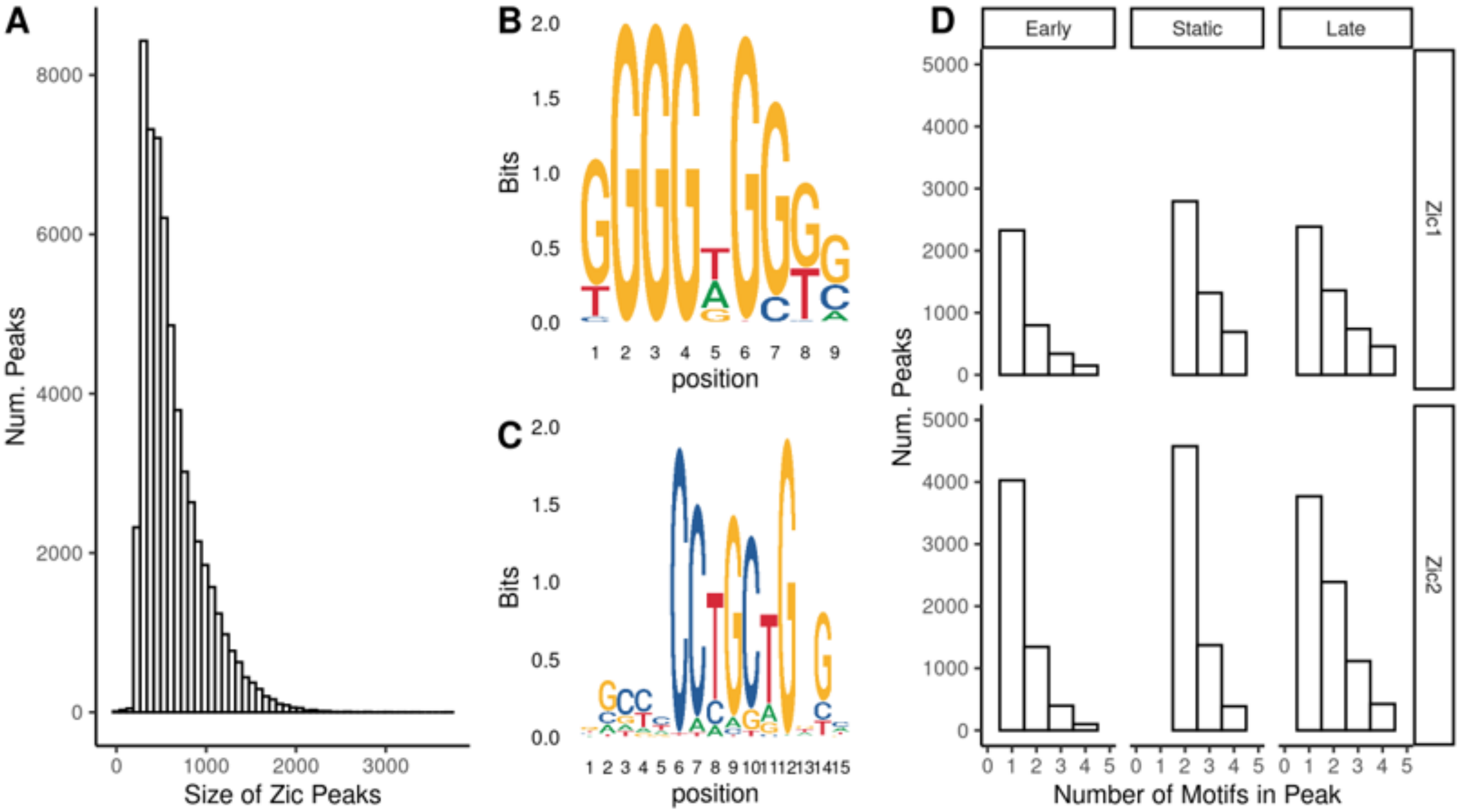

Supplementary Figure 2

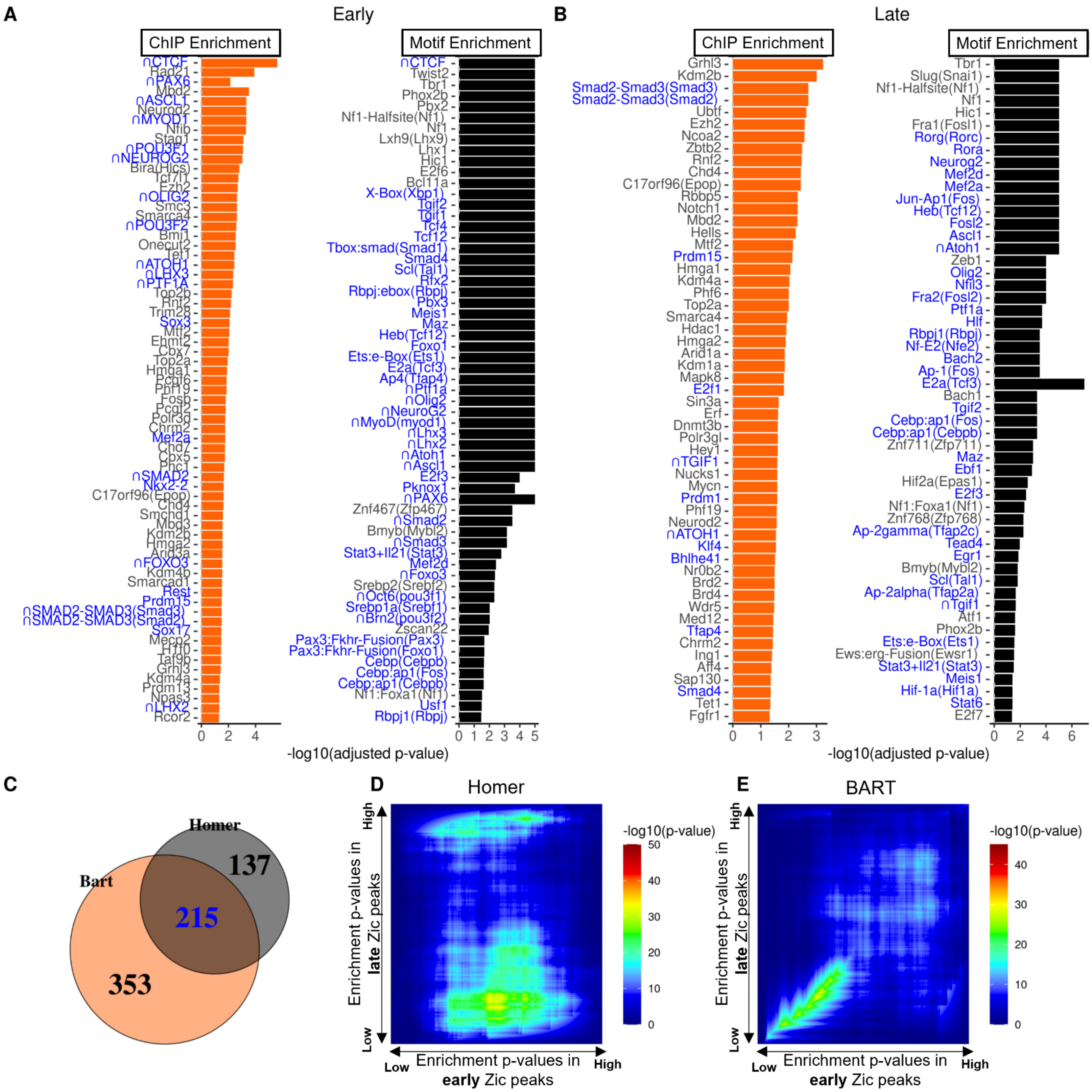

Supplementary Figure 3

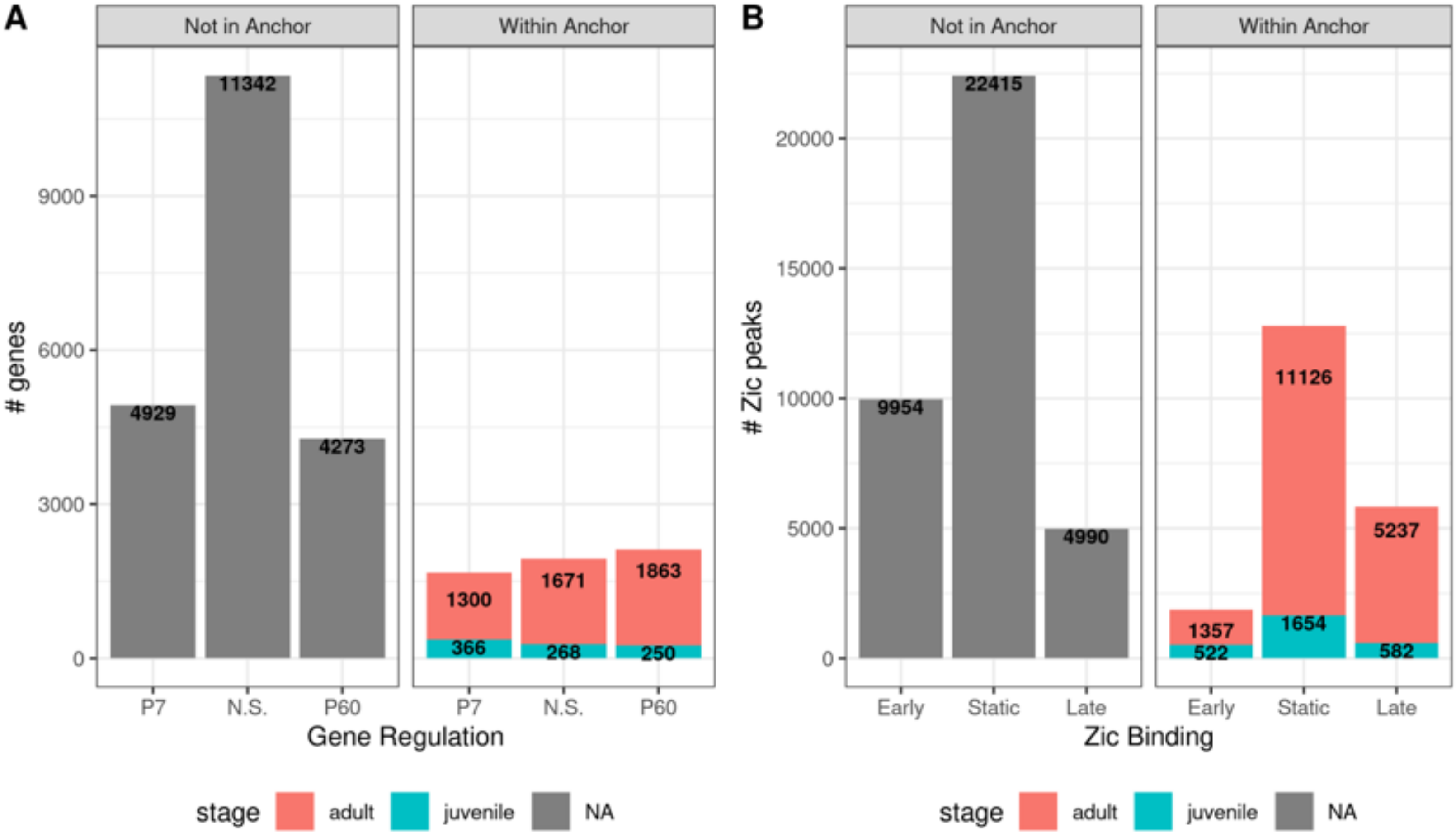

Supplementary Figure 4

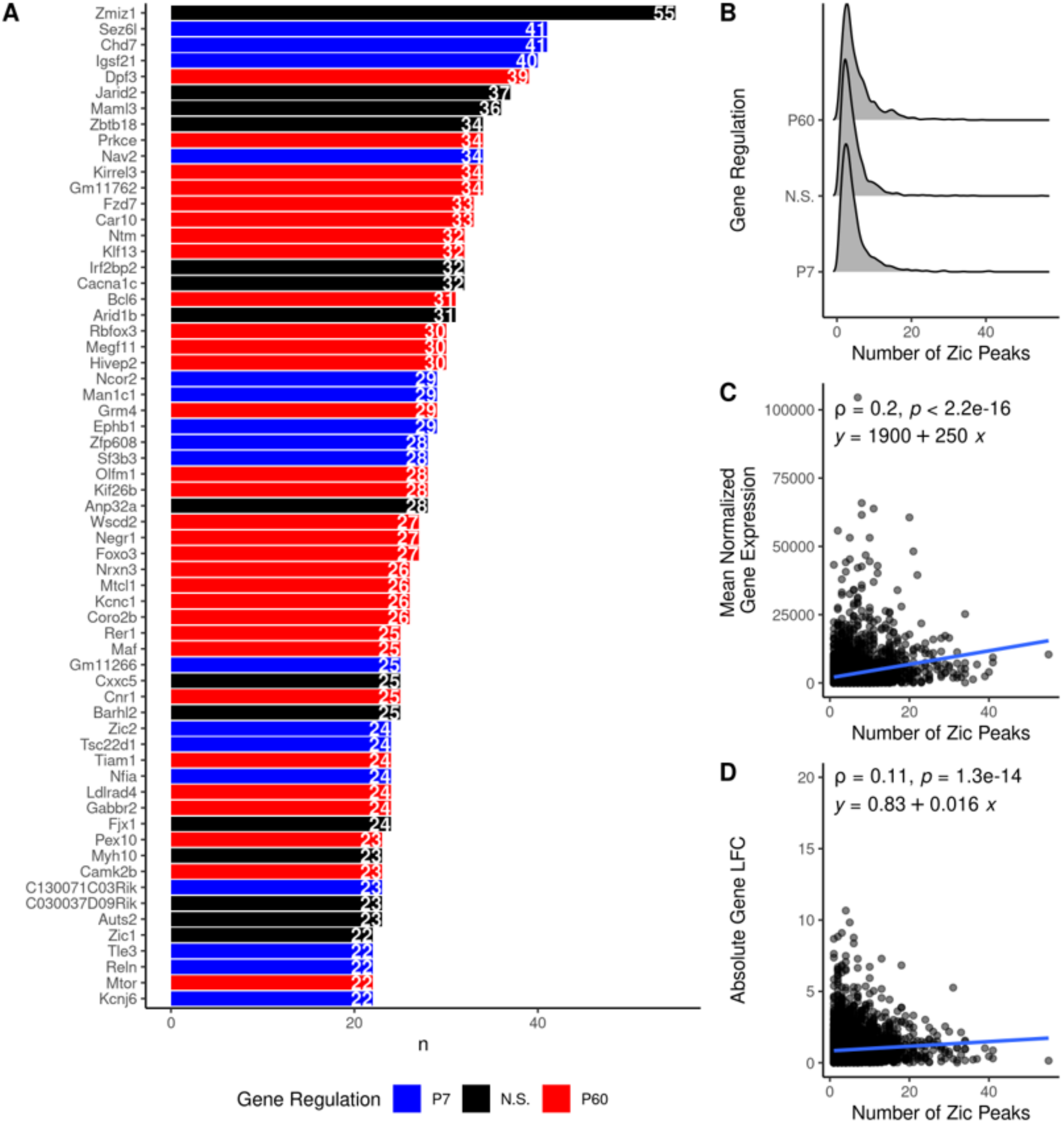

Supplementary Figure 5

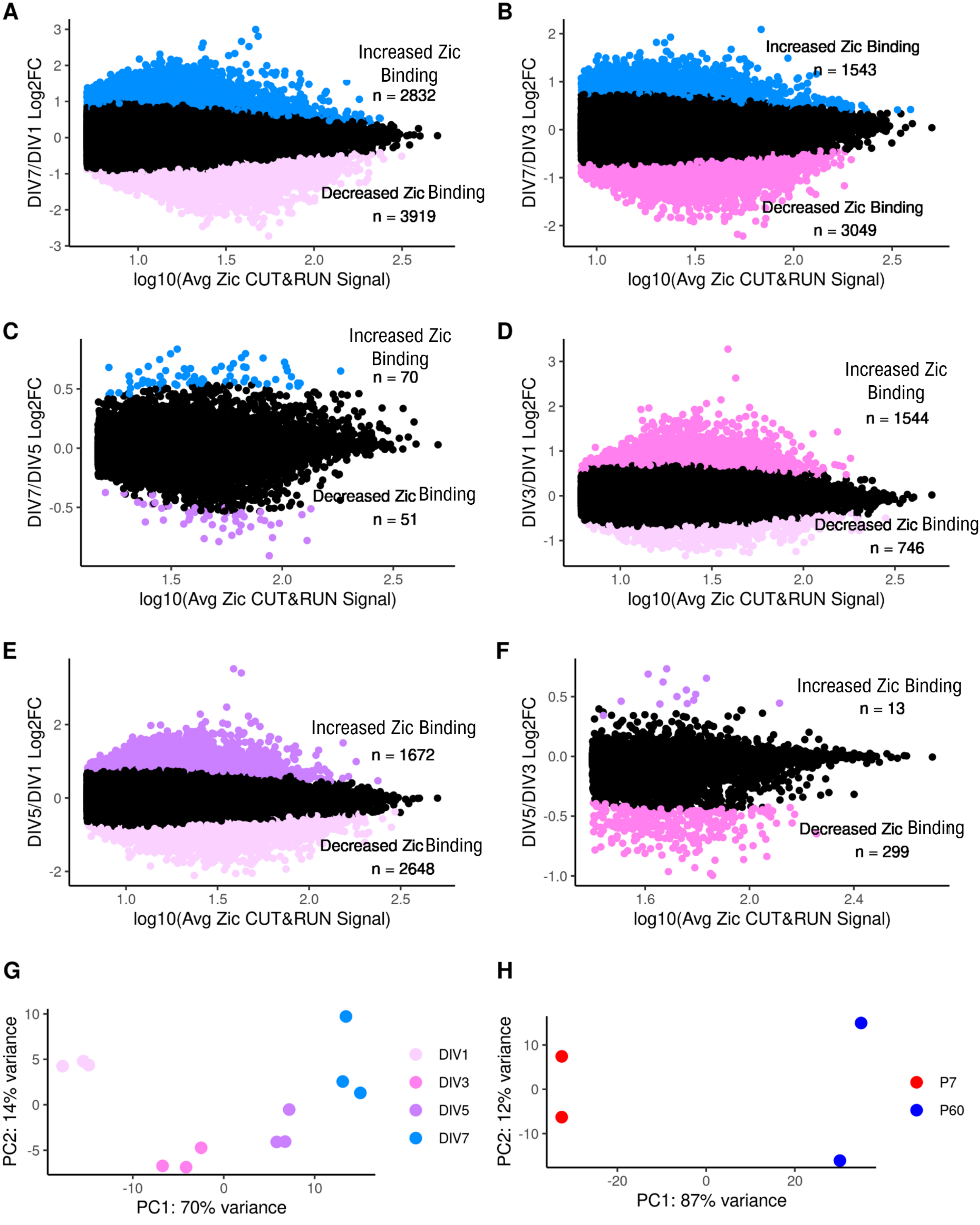

Supplementary Figure 6

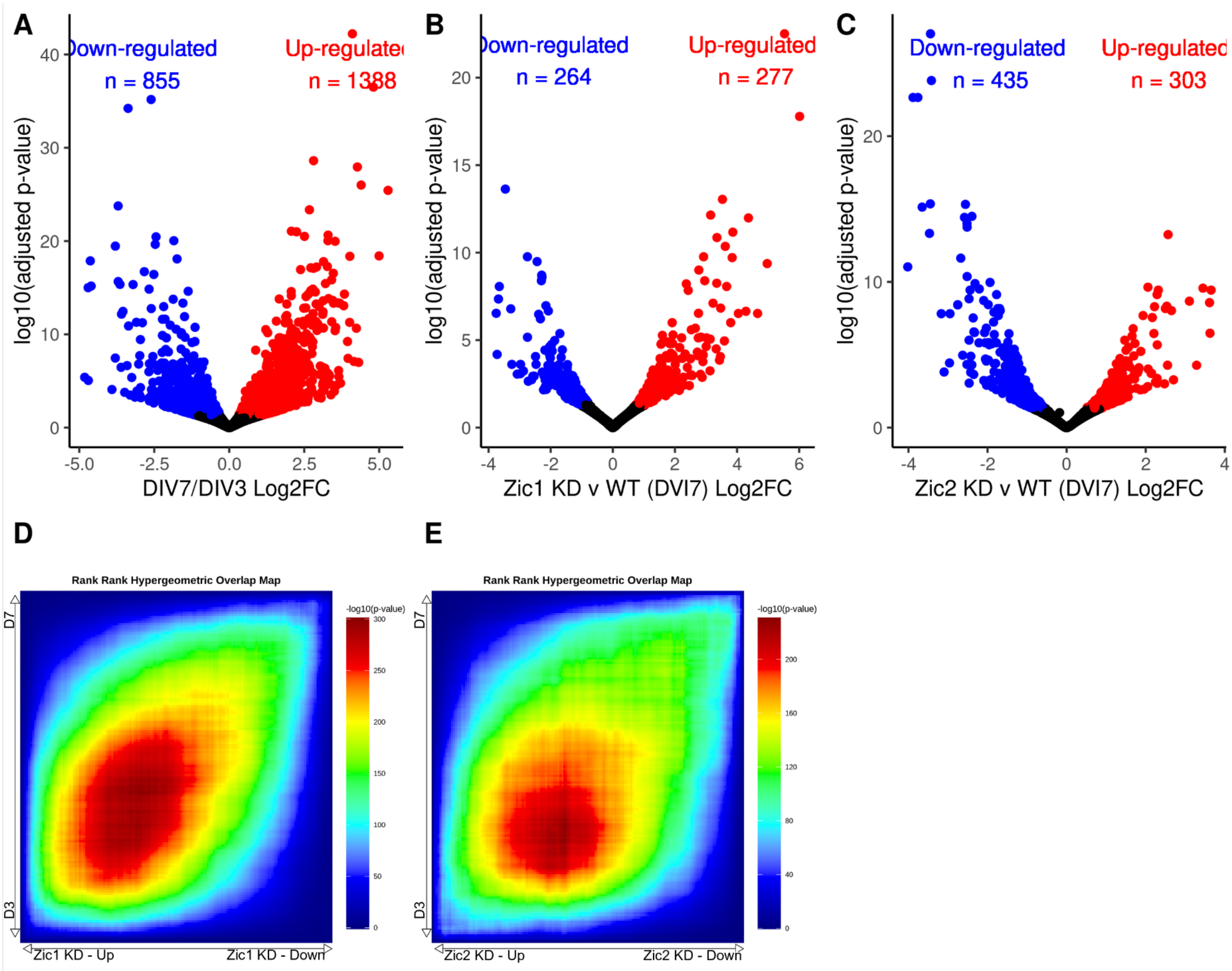

Supplementary Figure 7

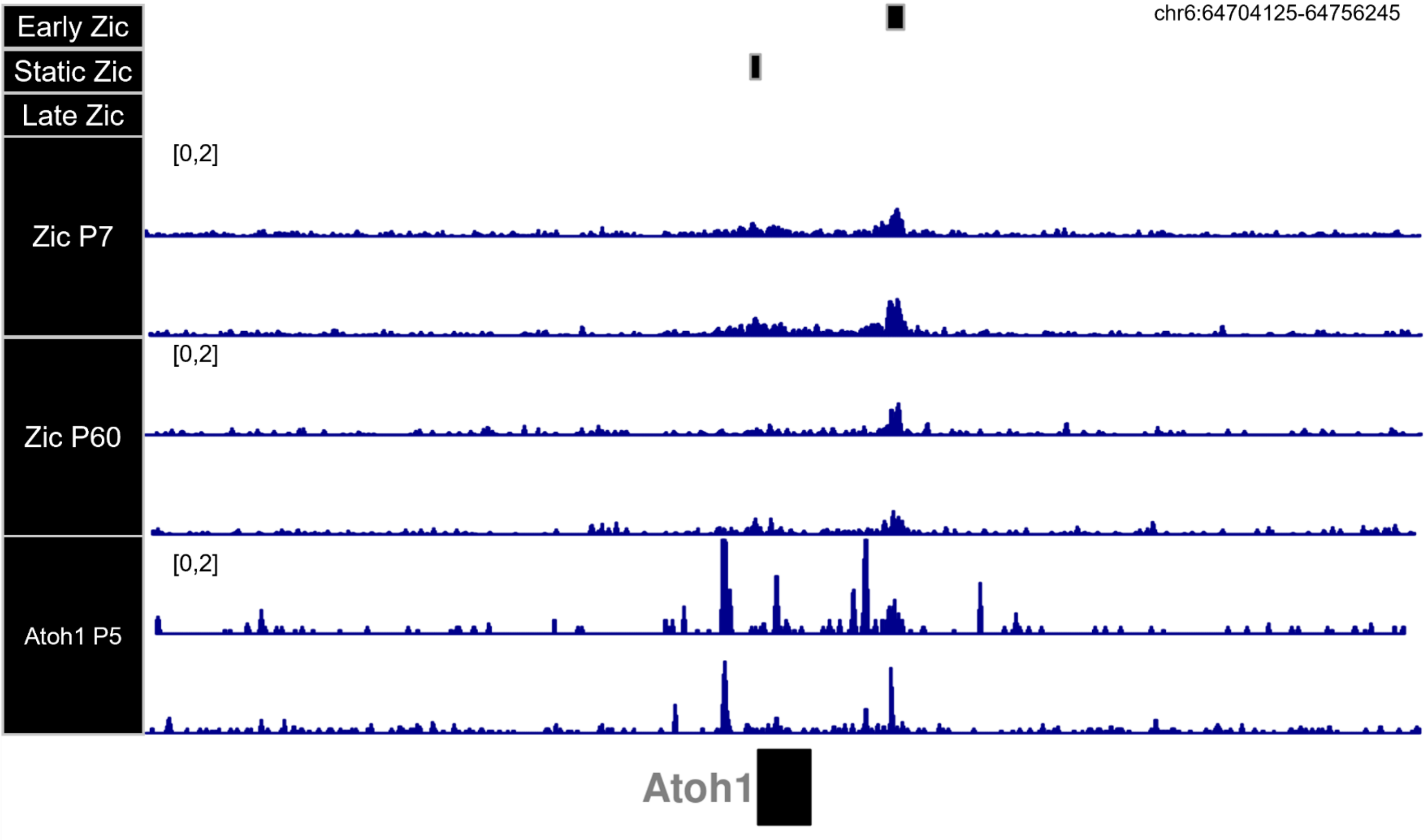
